# Loss of the fatty acid β-oxidation gene *acaa-2* impairs timely hatching under poor maternal diet in *C. elegans*

**DOI:** 10.64898/2026.09.29.755286

**Authors:** Yuzuha Komachiya, Riho Kato, Hayao Ohno

## Abstract

Developmental programs must remain robust despite fluctuations in maternal nutrition, yet the genetic basis of this robustness is incompletely understood. Using an unbiased, diet-dependent forward genetic screen in the nematode *Caenorhabditis elegans*, we searched for mutants with normal embryogenesis on standard food but defective development when mothers experienced poor nutrition. We isolated a mutation in *acaa-2*, which encodes an enzyme catalyzing the final step of mitochondrial fatty acid β-oxidation, and confirmed its role using an independent deletion allele. *acaa-2* mutant embryos exhibit a marked hatching delay specifically when mothers consume a low-quality diet. Despite remaining motile within the eggshell, mutant embryos are associated with increased unoccupied intra-eggshell space, consistent with impaired late embryonic growth. Our findings reveal a previously unrecognized role for *acaa-2* and suggest that mitochondrial fatty acid β-oxidation acts as a metabolic buffer that promotes developmental robustness under maternal nutrient limitation.

## Introduction

Embryogenesis in multicellular organisms is remarkably robust: developmental programs tend to proceed reliably even in the face of genetic variation or environmental perturbations (Félix & Barkoulas, 2015; Gilbert & Epel, 2015; Waddington, 1942). Such robustness is critical for successful survival and reproduction in the wild, and identifying its molecular underpinnings remains a major goal of developmental biology. However, much of the field’s work has been performed in artificially controlled laboratory settings where temperature, diet, and microbial exposure are regulated, leaving open the question of how embryogenesis is preserved under the kinds of environmental stresses—such as maternal nutrient limitation—that organisms commonly experience in nature (Félix & Duveau, 2012).

One of the principal metabolic strategies organisms employ to withstand environmental nutrient shortage is fatty acid β-oxidation (Bartlett & Eaton, 2004; Houten & Wanders, 2010; Poirier et al., 2006). This pathway, operating in mitochondria and peroxisomes, degrades acyl-CoA stepwise to yield acetyl-CoA and NADH and FADH_2_, which fuel ATP production. The final step of the cycle is mediated by thiolase I, also known as acetyl-CoA C-acyltransferase or 3-ketoacyl-CoA thiolase, a key enzyme for extracting energy from lipids (Bartlett & Eaton, 2004; Houten & Wanders, 2010). Previous studies in mammals have shown that fatty acid β-oxidation can function both as an energy source and as a regulator of signaling during early development and in the maintenance of tissue stem cells (Dunning et al., 2014; Ito et al., 2012; Knobloch et al., 2017).

The nematode *Caenorhabditis elegans*, with its fully mapped cell lineage and facile genetic manipulability (Brenner, 1974; Sulston et al., 1983), has long served as a powerful model for dissecting links between environmental cues and development. Fatty acid β-oxidation underlies key responses to nutrient stress in *C. elegans*—including starvation responses, dauer formation, and lifespan modulation—and lipid metabolism regulated through the nuclear receptor NHR-49 contributes to starvation resistance and metabolic adaptation (Butcher et al., 2009; Park & Paik, 2017; Pathare et al., 2012; Ratnappan et al., 2014; Van Gilst, Hadjivassiliou, Jolly, et al., 2005; Van Gilst, Hadjivassiliou, & Yamamoto, 2005). However, most investigations have focused on larval or adult physiology, and which metabolic pathways embryos use to complete development when maternal nutrition is limited remains poorly understood.

Across animal species, maternal environment can shape offspring development and physiology. In *C. elegans*, maternal feeding conditions alter traits including egg size, yolk accumulation, stress resistance, larval growth rate, and alternative embryogenesis associated with changes in somatic cell number (Hibshman et al., 2016; Jordan et al., 2019; Ohno & Bao, 2022). Early embryogenesis is highly dependent on maternal mRNAs and proteins deposited during oogenesis; their translation, activation, and degradation are tightly controlled after fertilization (Robertson & Lin, 2015; Stitzel & Seydoux, 2007).

The nutritional status of *C. elegans* is strongly influenced by the species of bacteria it eats. For example, *Bacillus megaterium* (also known as *Priestia megaterium*) is substantially larger than standard *Escherichia coli* strains such as OP50 or HB101 and is therefore less readily consumed by the worm; feeding on *B. megaterium* induces growth retardation and a state resembling dietary restriction (Avery & Shtonda, 2003; Shtonda & Avery, 2006). Recent studies have further demonstrated that the choice of bacterial diet can markedly alter nematode metabolism, reproductive output, lifespan, and other physiological traits (Samuel et al., 2016; Stuhr & Curran, 2020; Watson et al., 2014). Nevertheless, the molecular mechanisms that allow embryos to complete development when mothers experience poor nutrition remain largely unknown. A productive approach to identifying factors that confer developmental robustness is to screen for mutants that exhibit embryonic defects only under environmental stress.

In this study, we performed a forward genetic screen in *C. elegans* to identify mutants whose embryogenesis is abnormal only when mothers experience a low-quality bacterial diet. From the screen, we isolated a mutation in *acaa-2*, a *C. elegans* ortholog of thiolase I that catalyzes the terminal step of mitochondrial fatty acid β-oxidation. *acaa-2* mutants develop normally on standard food but exhibit hatching delays when their mothers are fed *B. megaterium*. The phenotype appears specific to the mitochondrial β-oxidation pathway and may reflect a late-embryonic growth defect rather than reduced embryonic motility.

## Results

To identify molecules that allow *C. elegans* embryos to progress normally under poor maternal nutrition, we screened for mutants whose embryogenesis is abnormal only when mothers are fed a low-quality diet (Fig. 1A). We mutagenized wild-type N2 animals with ethyl methanesulfonate (EMS), reared progeny on the high-quality food *E. coli* HB101 to the F_2_ generation, and then isolated individual F_2_ animals. F_3_ eggs were collected from each isolated F_2_ by brief exposure to sodium hypochlorite, and the resulting F_3_ embryos were divided and plated onto HB101 or onto the low-quality diet *B. megaterium* (DA1880). Although the precise nutritional differences available to the worm are not defined, DA1880 is commonly used to model dietary restriction in *C. elegans* because its large cells are less readily ingested, which slows worm growth (Avery & Shtonda, 2003; Shtonda & Avery, 2006). We inspected plates the day after F_3_ animals began egg-laying and selected lines in which F_4_ embryos hatched normally on HB101 but exhibited developmental abnormalities on DA1880.

**Figure 1.**
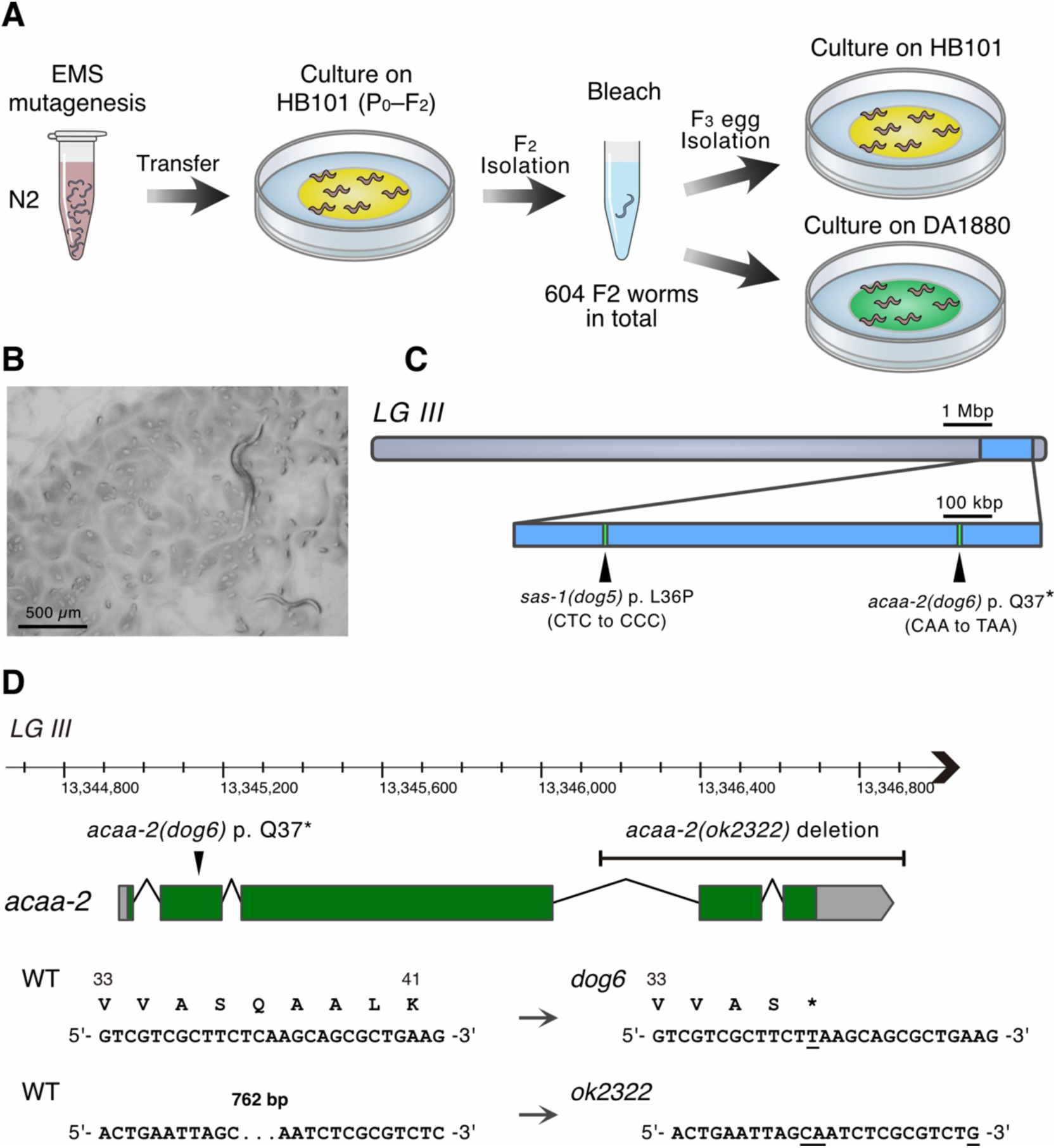
Forward genetic screen for diet-dependent embryogenesis defects. (**A**) Schematic of the genetic screen. F_3_ embryos were isolated from single F_2_ animals derived from EMS-mutagenized wild-type *C. elegans*, and matched cohorts of embryos were cultured on *E. coli* HB101 or *B. megaterium* DA1880 to assess F_4_ development. A total of 604 mutagenized lines were screened. (**B**) Representative photograph of mutagenized line No. 469 on a DA1880 plate. (**C**) Chromosomal region identified by repeated outcrossing to wild type and whole-genome sequencing. (**D**) Genomic organization of the *acaa-2* locus and the nucleotide-level and predicted protein-level changes caused by *acaa-2(dog6)* and *acaa-2(ok2322)*. *dog6* was isolated in this study; *ok2322* was obtained from the *C. elegans* Gene Knockout Consortium.

We conducted the screen on 604 lines derived from individual F_2_ animals. In 603 of the 604 lines (99.8%), we did not recover any mutant candidates that exhibited developmental defects only when mothers were fed DA1880, suggesting that embryogenesis shows robustness to the environmental variation tested. However, one exception, line No. 469, displayed a diet-dependent phenotype: embryos developed indistinguishably from wild type on HB101 but showed marked egg accumulation on DA1880 plates (Fig. 1B). Although most of these eggs eventually hatched on DA1880, hatching was markedly delayed compared with HB101.

To pinpoint the causative lesion, we applied the strategy of Zuryn et al. (2010): repeated outcrossing to the parental wild-type strain followed by whole-genome sequencing to identify genomic regions enriched for the characteristic EMS G/C > A/T transitions. Using the DA1880-dependent egg-accumulation phenotype, we independently generated two lines that had been outcrossed four times to wild type and subjected them both to whole-genome sequencing. The defect mapped to the right arm of chromosome III, and within this interval only two coding changes were identified: *sas-1(dog5)* and *acaa-2(dog6)* (Fig. 1C, D). *sas-1(dog5)* results in a p.Leu36Pro substitution in SAS-1, whereas *acaa-2(dog6)* introduces a premature stop codon at Gln37 (p.Gln37*) that truncates ACAA-2 (Fig. 1C, D). After additional crosses that separated the two lesions, we obtained isolates carrying each mutation individually; the diet-dependent hatching delay persisted only in the *acaa-2(dog6)* single mutant (Fig. 2A, B). We also analyzed *acaa-2(ok2322)*, a deletion allele generated by the *C. elegans* Deletion Mutant Consortium (*C. elegans* Deletion Mutant Consortium, 2012). PCR and Sanger sequencing showed that *ok2322* is a deletion that removes exons 4 and 5 of *acaa-2* (Fig. 1D). After five backcrosses into the N2 background, the *ok2322* strain displayed the same pronounced delay in hatching on DA1880 as *acaa-2(dog6)* (Fig. 2A, B). The concordant, diet-dependent phenotype of two independent, outcrossed alleles strongly implicates *acaa-2* as the causal locus. A complementation test further confirmed this: embryos from mothers carrying the two alleles in trans (*acaa-2(dog6)*/*acaa-2(ok2322)* trans-heterozygotes) displayed the same DA1880 hatching defect as *acaa-2* homozygotes (Fig. 2C).

**Figure 2.**
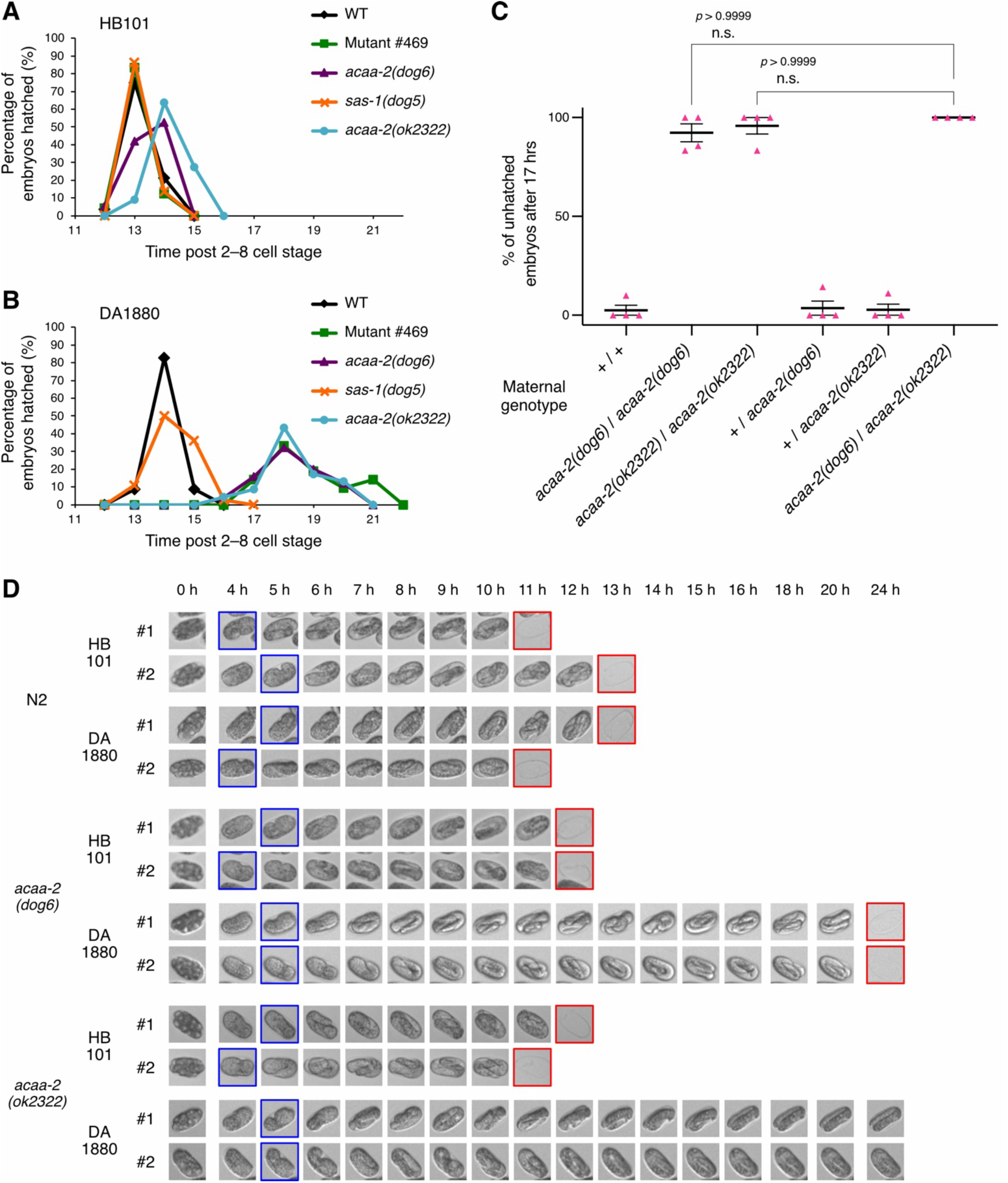
Hatching is delayed in *acaa-2* mutant embryos when mothers are fed DA1880. (**A**, **B**) Wild type, mutagenized line No. 469, *acaa-2(dog6)*, *sas-1(dog5)*, and *acaa-2(ok2322)* were reared on HB101 (A) or DA1880 (B). Early-stage embryos were isolated and time to hatching was recorded. n = 19– 36 embryos. (**C**) Proportion of embryos that remained unhatched 17 hours after isolation from DA1880-reared mothers for: wild type (+/+), *acaa-2(dog6)* homozygotes, *acaa-2(ok2322)* homozygotes, *acaa-2(dog6)* heterozygotes, *acaa-2(ok2322)* heterozygotes, and *acaa-2(dog6)*/*acaa-2(ok2322)* trans-heterozygotes. n = 4 independent assays (32–36 embryos total). Kruskal-Wallis and Dunn’s post-test. n.s., not significant. (**D**) Representative 24-h time-lapse images of early embryos isolated into buffer after maternal rearing on HB101 or DA1880; blue boxes indicate the comma stage and red boxes indicate hatching.

To determine when *acaa-2* mutant embryos become delayed under poor maternal nutrition, we isolated *acaa-2(dog6)* and *acaa-2(ok2322)* embryos into buffer and performed long-term time-lapse imaging with a compound microscope (Fig. 2D). Both alleles progressed to the comma stage with little apparent delay; however, the subsequent elongation and three-fold stages were substantially prolonged in embryos whose mothers had been reared on DA1880 compared with wild type (Fig. 2D). These observations may indicate that *acaa-2* function is particularly important during late embryogenesis when maternal diet quality is low.

ACAA-2 is an ortholog of human acetyl-CoA acyltransferase 2 (ACAA2), a member of the thiolase I family. ACAA-2 and hydroxyacyl-CoA dehydrogenase trifunctional multienzyme complex subunit beta (HADHB) catalyze the terminal step of mitochondrial fatty acid β-oxidation (Fig. 3A) and are therefore thought to be important for mobilizing lipids to generate ATP. We tested putative loss-of-function mutants of other worm thiolase homologs — *hadb-1*, *kat-1*, *daf-22* and *T02G5.7* (Fig. 3B) — alongside *acaa-2(dog6)* and *acaa-2(ok2322)* for diet-dependent hatching defects. The *hadb-1(tm6853)* mutant showed an abnormal hatching phenotype when mothers were reared on DA1880, although the difference relative to HB101 was not statistically significant (Fig. 3C). By contrast, mutants affecting the peroxisomal β-oxidation factor *daf-22* or the thiolase II homologs *kat-1* and *T02G5.7* showed no difference in hatching between HB101 and DA1880 conditions (Fig. 3C). Together, these results suggest that mitochondrial fatty acid β-oxidation is particularly important for embryos to complete development at normal speed when maternal diet quality is low.

**Figure 3.**
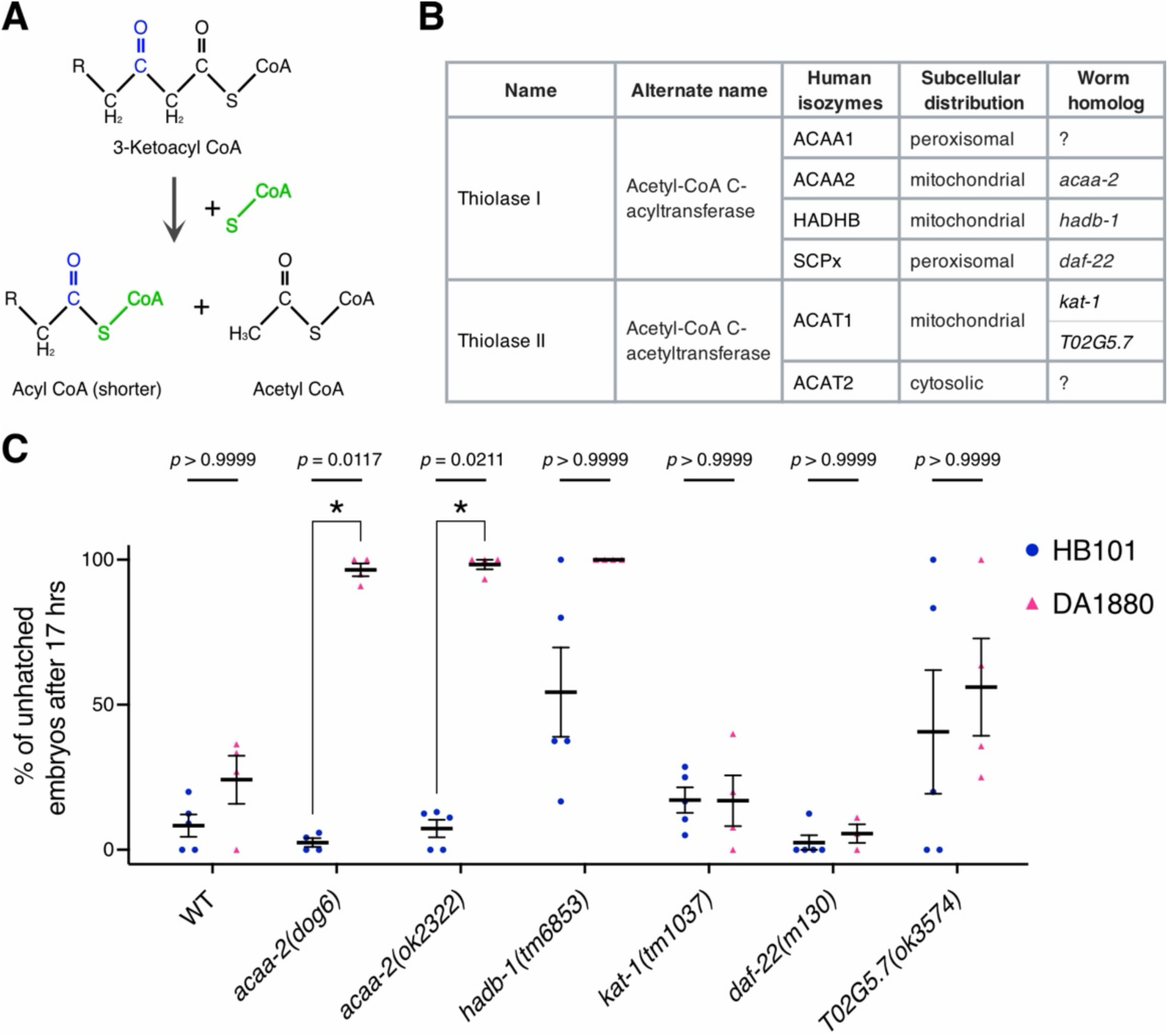
Mitochondrial β-oxidation contributes to completion of embryogenesis when mothers are reared on DA1880. (**A**) Schematic of the reaction catalyzed by thiolase I. (**B**) Human thiolase enzymes and their putative *C. elegans* homolog genes. (**C**) Mothers carrying thiolase mutations were reared on HB101 or DA1880; early embryos were isolated, incubated in buffer, and the proportion remaining unhatched after 17 h was recorded. n = 3–5 independent assays (48–83 embryos total). Kruskal-Wallis and Dunn’s post-test.

To determine whether *acaa-2* acts in the mother or in the embryo, we mated DA1880-reared *acaa-2(dog6)* hermaphrodites to wild-type males, producing cross-progeny that inherited the wild-type *acaa-2* allele but were born from *acaa-2* mutant mothers. These cross-progeny did not show delayed hatching on DA1880 (Fig. 4A), implying that zygotic *acaa-2* expression is sufficient to rescue on-time embryogenesis and that *acaa-2* can act in the embryo under poor maternal diet. However, embryos derived from mothers heterozygous for *acaa-2* likewise hatched without delay on DA1880 (Fig. 2C). To distinguish the genotypes of embryos produced by heterozygous mothers, we generated a strain heterozygous for *acaa-2(dog6)* and carrying the fluorescently marked balancer chromosome *tmC29*[*tmIs1259*] (Dejima et al., 2018) and examined development of *acaa-2(dog6)* homozygous embryos identified by absence of balancer fluorescence. These homozygous embryos also lacked the DA1880-dependent hatching delay (Fig. 4B), consistent with a maternal contribution. Therefore, maternally provided *acaa-2* transcripts or protein may persist in the embryo, or a substance produced by maternal ACAA-2 activity may be transferred to and function within the embryo.

**Figure 4.**
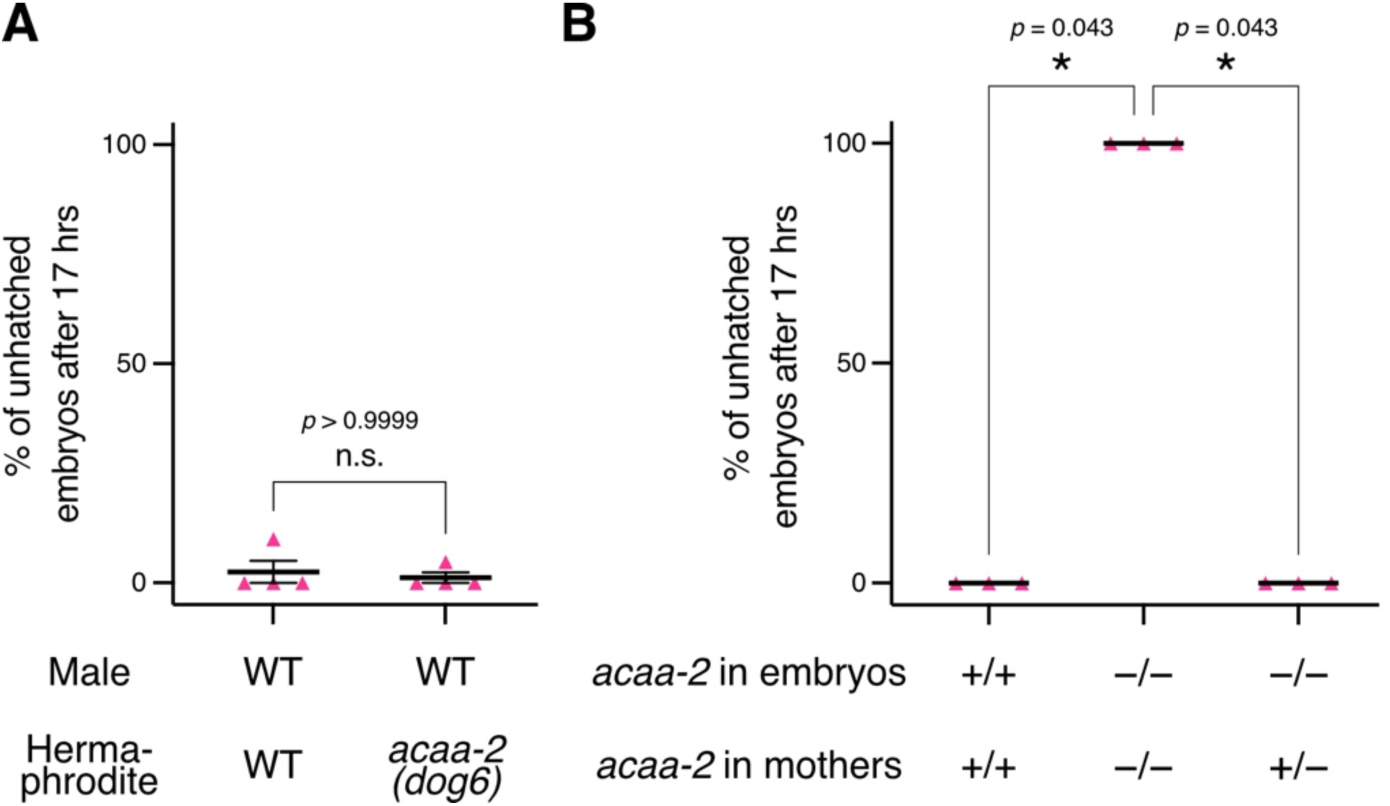
Zygotic or maternal *acaa-2* expression rescues the DA1880-dependent hatching defect. (**A**) Wild-type males were mated to either wild-type or *acaa-2(dog6)* hermaphrodites on DA1880; isolated embryos derived from cross-fertilization were scored for the fraction remaining unhatched after 17 h. n = 4 independent assays (26–44 embryos in total). Two-tailed Mann-Whitney test. (**B**) Proportion of embryos remaining unhatched 17 h after isolation for embryos from wild-type mothers, embryos from *acaa-2(dog6)* homozygous mothers, and *acaa-2(dog6)* homozygous embryos produced by *acaa-2(dog6)* heterozygous mothers (*acaa-2(dog6)*/*tmC29*[*tmIs1259*]). n = 3 independent assays (18–67 embryos in total). Kruskal-Wallis and Dunn’s post-test.

To investigate the basis of the delayed hatching in DA1880-fed *acaa-2* mutants, we considered whether loss of ACAA-2–dependent β-oxidation might diminish the embryo’s capacity for the forceful movements required for eggshell rupture. We therefore measured motility in three-fold stage embryos. Contrary to this energy-deficit hypothesis, *acaa-2* mutant embryos from DA1880-reared mothers moved more vigorously inside the eggshell than wild-type embryos did (Fig. 5A, Movie 1–4). One plausible explanation for this apparent hyperactivity is that the embryo has more room to move: although eggshell area was not increased in DA1880-reared *acaa-2* mutants (Fig. 5B), the blank space not occupied by the embryo was larger than in either DA1880-reared wild type or HB101-reared *acaa-2* mutants (Fig. 5C, D, Movie 1–4). These observations raise the possibility that *acaa-2* mutant embryos fail to grow sufficiently within the eggshell under poor maternal diet, and that this lack of internal volume reduces the mechanical pressure available to break the eggshell at hatching.

**Figure 5.**
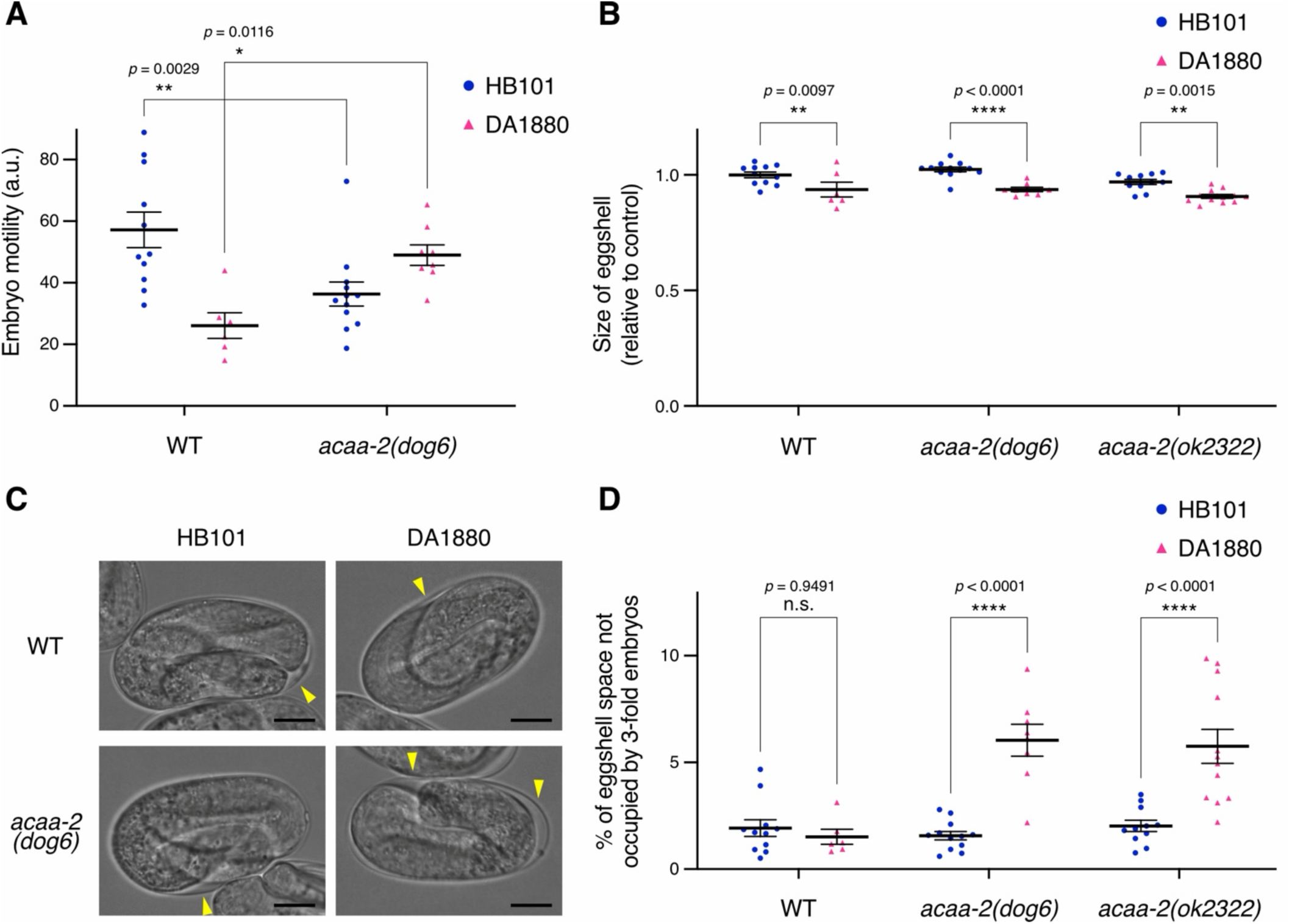
Embryos from DA1880-fed mothers show an abnormal size, not reduced motility. (**A**) Quantification of embryonic motility at the three-fold stage for wild type and *acaa-2(dog6)* embryos whose mothers were reared on HB101 or DA1880. Early embryos were isolated and incubated for 8.5 h to reach the three-fold stage prior to imaging; motility is reported in arbitrary units (a.u.). n = 6–12 embryos. ANOVA with Šídák’s post hoc test. (**B**) Eggshell area measured from micrographs of the indicated genotypes reared on HB101 or DA1880. n = 6–12 embryos. ANOVA with Šídák’s post hoc test. (**C**) Representative bright-field images of three-fold stage embryos; yellow arrowheads indicate the unoccupied intra-eggshell space. Scale bar = 10 µm. (**D**) Quantification of the proportion of eggshell area unoccupied by the embryo. n = 6–12 embryos. ANOVA with Šídák’s post hoc test.

## Discussion

In this study, we screened for mutants that display embryonic defects only when mothers are reared on the low-quality bacterial diet *B. megaterium* and identified *acaa-2*, which encodes the mitochondrial thiolase I of the fatty acid β-oxidation pathway, as the causal locus. Under standard *E. coli* feeding, *acaa-2* mutants show little or no developmental abnormality, but when mothers consume *B. megaterium*, they show marked delays in hatching due to defects that manifest late in embryogenesis. The phenotype was not observed for mutants in the peroxisomal β-oxidation gene *daf-22* or in thiolase II homologs, indicating a relative specificity for the mitochondrial β-oxidation branch. The delayed hatching does not correlate with reduced embryonic motility; rather, it may be associated with a late-embryonic growth defect in which embryos fail to occupy the eggshell fully. Together, these findings suggest that *acaa-2*–dependent fatty acid β-oxidation provides a critical metabolic buffer that contributes to the robustness of embryogenesis when maternal diet quality is poor.

One notable outcome of this study is the first report, to our knowledge, of an *acaa-2*– dependent phenotype in *C. elegans*. Prior studies of fatty acid β-oxidation in the worm have largely emphasized NHR-49–mediated metabolic regulation, starvation responses, dauer formation, and lifespan control (Ratnappan et al., 2014; Van Gilst, Hadjivassiliou, Jolly, et al., 2005; Van Gilst, Hadjivassiliou, & Yamamoto, 2005), whereas the embryonic phenotypes of β-oxidation mutants have not been thoroughly characterized. Both *acaa-2* alleles we examined developed normally when reared on standard *E. coli*, implying that conventional laboratory diets may mask *acaa-2*’s developmental role. Our results therefore illustrate how screening for environment-specific phenotypes can reveal mechanisms of developmental robustness that remain hidden under typical culture conditions.

Our results also suggest that fatty acid β-oxidation can support embryogenesis in a manner that depends on maternal state. Lipids are well suited for long-term energy storage, and the mobilization of those stores by β-oxidation during nutrient scarcity is widely recognized as important for organismal survival (Bartlett & Eaton, 2004; Houten & Wanders, 2010). Nevertheless, to our knowledge, there are relatively few reports that directly link disruption of the β-oxidation pathway to delayed or failed embryogenesis. Our findings are consistent with a model in which fatty acid β-oxidation in *C. elegans* does not have a uniform, constitutive role in embryogenesis but instead acts as a metabolic buffer that helps preserve developmental robustness when maternal diet quality is low.

Nonetheless, the precise biochemical cause of the *acaa-2*–dependent hatching delay is still unknown. A simple and plausible possibility is that impaired mitochondrial β-oxidation lowers ATP production, compromising late embryonic growth. Alternatively, the pathway’s product, acetyl-CoA, might be required for chromatin modifications such as histone acetylation (McDonnell et al., 2016) that regulate developmental gene expression. Other possibilities include effects of metabolic water generated during fatty-acid breakdown or developmental defects caused by accumulation or depletion of fatty acids of particular chain lengths or their derivatives. It is also noteworthy that *hadb-1* mutants showed a phenotype of different severity than *acaa-2* mutants (Fig. 3C). In mammals, HADHB functions primarily in long-chain fatty-acid degradation, whereas ACAA2 acts on medium- to short-chain substrates (Houten & Wanders, 2010); if *C. elegan*s shares this substrate partitioning, loss of *hadb-1* might deprive ACAA-2 of its substrates and thereby produce a stronger effect than loss of *acaa-2* itself. Further work—such as lipid profiling and metabolic-flux analyses—will be required to resolve how these enzymes functionally interact.

There are important limitations of this study to consider. First, we did not profile maternal or embryonic lipid composition under HB101 and DA1880 conditions, leaving unclear why reliance on lipid catabolism increases under DA1880. Second, although *acaa-2* mutant embryos from DA1880-reared mothers show an enlarged unoccupied space inside the eggshell, we have not demonstrated that this enlargement is the direct cause of the hatching delay. Third, our genetic data indicate that zygotic *acaa-2* expression is sufficient for phenotypic rescue, yet embryos produced by heterozygous mothers are also protected, suggesting a maternal contribution. Whether maternally deposited transcripts or proteins persist in the embryo, or whether a metabolite synthesized by maternal ACAA-2 is transferred to the embryo, remains to be determined. Addressing these issues in future studies will help reveal the metabolic strategies that preserve embryonic robustness in response to changes in maternal nutrition.

## Materials and Methods

### Strains and culture

*C. elegans* strains were maintained at 20°C following standard methods (Brenner, 1974). Animals were cultured on nematode growth medium (NGM) plates seeded with either *E. coli* HB101 or *B. megaterium* DA1880. Bristol N2 was used as the wild-type strain. All strains and oligonucleotide primers used in this study appear in Tables S1 and S2, respectively.

### Genetic screen and whole-genome sequencing

EMS mutagenesis was carried out as described by Brenner (1974). Approximately 10,000 L4 N2 animals were mutagenized and maintained as 20 separate populations until the F_2_ generation. When F_2_ animals reached adulthood, approximately 30 individuals from each population (604 animals total) were picked singly into individual wells of a 96-well plate. Prior to adding worms, Clorox™ disinfecting bleach was diluted 1:5 (v/v) in water; 1 µL of this diluted solution was dispensed into each well and worms were then transferred into the wells. After a few minutes at room temperature, adult bodies were dissolved and intact fertilized F_3_ embryos remained. Each well received 20 µL M9 buffer (3 g/L KH_2_PO_4,_ 6 g/L Na_2_HPO_4_, 5 g/L NaCl, 1 mM MgSO_4_, 0.03% gelatin); the embryo suspensions were split, and half were plated onto NGM plates seeded with HB101 while the remainder were plated onto plates seeded with DA1880. Plates were examined on day 4 for HB101 and day 5 for DA1880; one line that produced many F4 embryos that hatched on HB101 but not on DA1880 was selected for further study.

To map the causal mutation, we followed Zuryn et al. (2010). The candidate line was outcrossed to N2 four times; two independent isolates derived from this outcrossing process were subjected to whole-genome sequencing on an Illumina NovaSeq platform to a mean coverage ≥20×. Sequence reads were analyzed with CloudMap (Minevich et al., 2012) to identify a genomic region enriched for EMS-signature G/C > A/T transitions and to extract candidate coding changes.

### Hatching assay

Early embryos were isolated as previously described (Ohno & Bao, 2022). Embryos at the 2– 30-cell stage were transferred either to unseeded NGM plates (Figs. 2A, 2B, 2C, 4A, 4B) or to individual wells of an 8-well chambered cover glass (Matsunami, SCC-008) containing 600 µL M9 buffer per well (Figs. 2D, 3C). After incubation at 20–22°C for 0–24 h (as indicated), hatching was scored using either a stereomicroscope (Evident SZX10; Figs. 2A, 2B, 2C, 4A, 4B) or the Axio Observer 7 inverted microscope system (Zeiss) equipped with an ORCA-Fusion BT camera (Hamamatsu) and Colibri 7 light source (Zeiss) (Figs. 2D, 3C). A UPLXAPO20X dry objective (Evident; NA = 0.80) was used with the Axio Observer 7 for scoring. For the experiments shown in Figs. 2C, 2D, 3C, 4A, and 4B, animals were reared on HB101 until the early L4 larval stage, then transferred to DA1880 plates; embryos were isolated 24–30 h after the transfer. In Fig. 4A, embryos that displayed Venus fluorescence in body-wall muscle at the three-fold stage (derived from mating with males carrying *Ex*[*myo-3^prom^::venus*]) were scored as cross-progeny. In Fig. 4B, embryos produced by mothers heterozygous for the *tmC29*[*tmIs1259*] balancer were scored as *acaa-2(dog6)* homozygotes if they lacked *tmIs1259*-derived green fluorescence at the three-fold stage.

### Microscopy and image analyses

For the experiments shown in Figs. 5A–5D and Movies 1–4, embryos at the 8–30-cell stage were isolated as described (Ohno & Bao, 2022), transferred to unseeded NGM plates, and incubated at 22°C for 8.5 h to reach the three-fold stage. As in Komachiya et al. (2026), a cut piece of NGM agar was inverted onto a 25 × 36 mm coverslip (Matsunami, Cat. No. C025361) and mounted on an inverted microscope (Axio Observer 7, Zeiss) equipped with an ORCA-Fusion BT camera (Hamamatsu Photonics) and Colibri 7 LED light source (Zeiss). Images were acquired with a PLFLN100X dry objective (Evident; NA = 0.95) in streaming mode for 10 s, yielding approximately 845 frames per acquisition (≈84.5 frames/s). Eggshell area and the area of the empty intra-eggshell space were measured on the single frame showing maximal unoccupied space using Fiji (Schindelin et al., 2012); regions were delineated with the polygon selection tool and areas recorded.

Embryo motility was quantified by frame-to-frame difference imaging in Fiji. For each time series a copy shifted by one frame (t = N → t = N+1) was created and subtracted from the original series using Image Calculator (Operation = Subtract) to produce difference images. For each difference image the embryo was outlined as a region of interest (ROI) and the mean pixel intensity within the ROI was measured. The motility metric for an embryo was calculated as the arithmetic mean of these ROI mean intensities across all difference frames. A nearby background ROI was measured in the same way and its mean (averaged across frames) was subtracted from the embryo mean to yield a background-corrected motility score.

### Statistical analysis

Statistical analyses were performed with a statistic package (Prism v.11.1.0, GraphPad software). Error bars indicate mean ± SEM. Statistical comparisons were performed with ANOVA with Šídák’s post hoc test, two-tailed Kruskal-Wallis with Dunn’s post-test, or two-tailed Mann-Whitney test.

## Acknowledgments

We thank the *Caenorhabditis* Genetics Center (CGC), which is funded by the NIH Office of Research Infrastructure Programs (P40OD010440), and the National Bioresource Project (NBRP), Japan, for providing strains. We also thank Dr. Zhirong Bao and members of the Bao lab for helpful discussions and suggestions, and the Integrated Genomics Operation (IGO) at Memorial Sloan Kettering Cancer Center for assistance with whole-genome sequencing. This work was supported by HFSP Long-Term Fellowship (LT000938/2017) and by JSPS KAKENHI Grant JP26K09212 awarded to H.O. The authors declare no competing interests. The raw data will be deposited on Figshare.

---

**Table S1:**
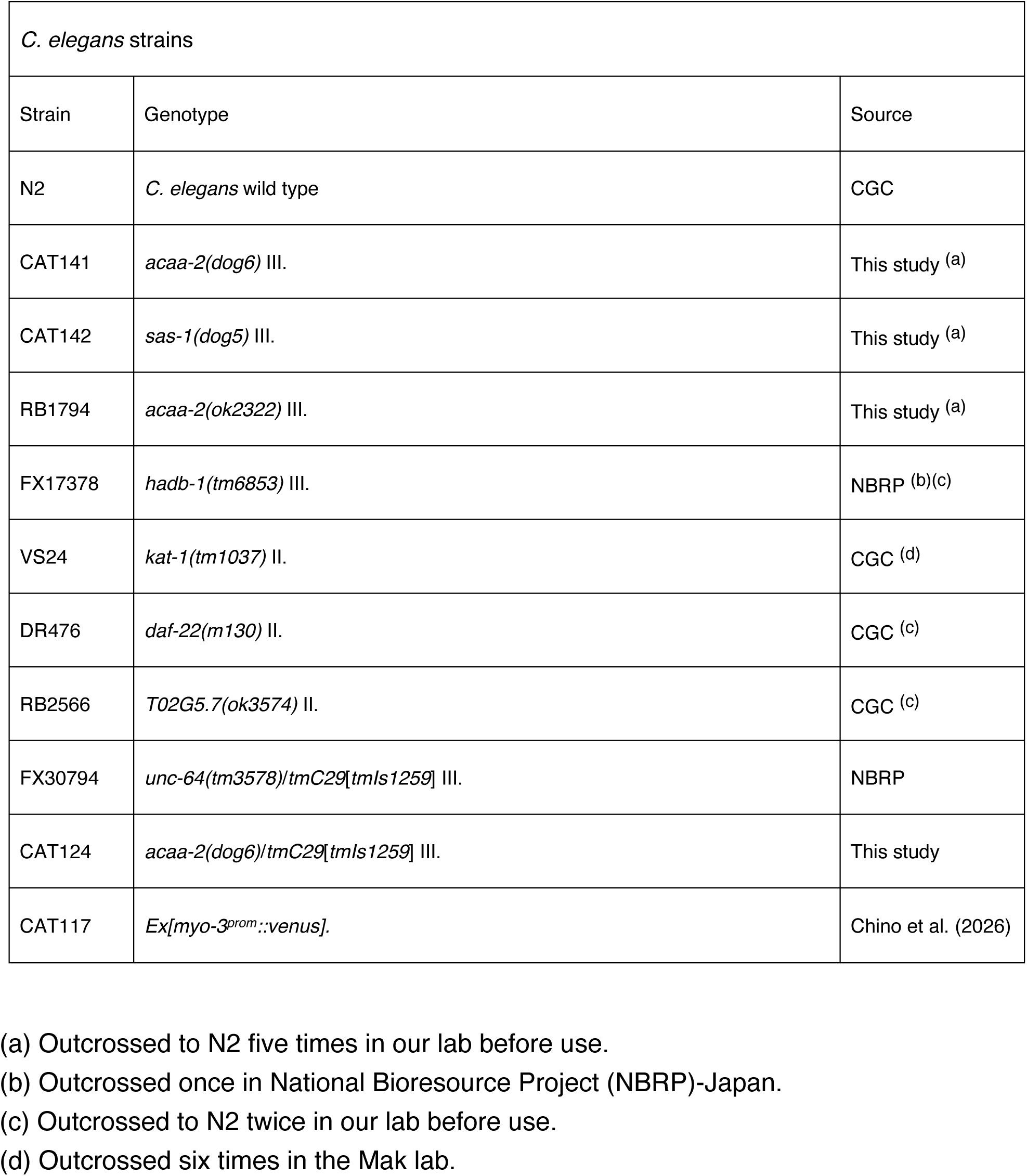
Strains used in this study.

**Table S2:**
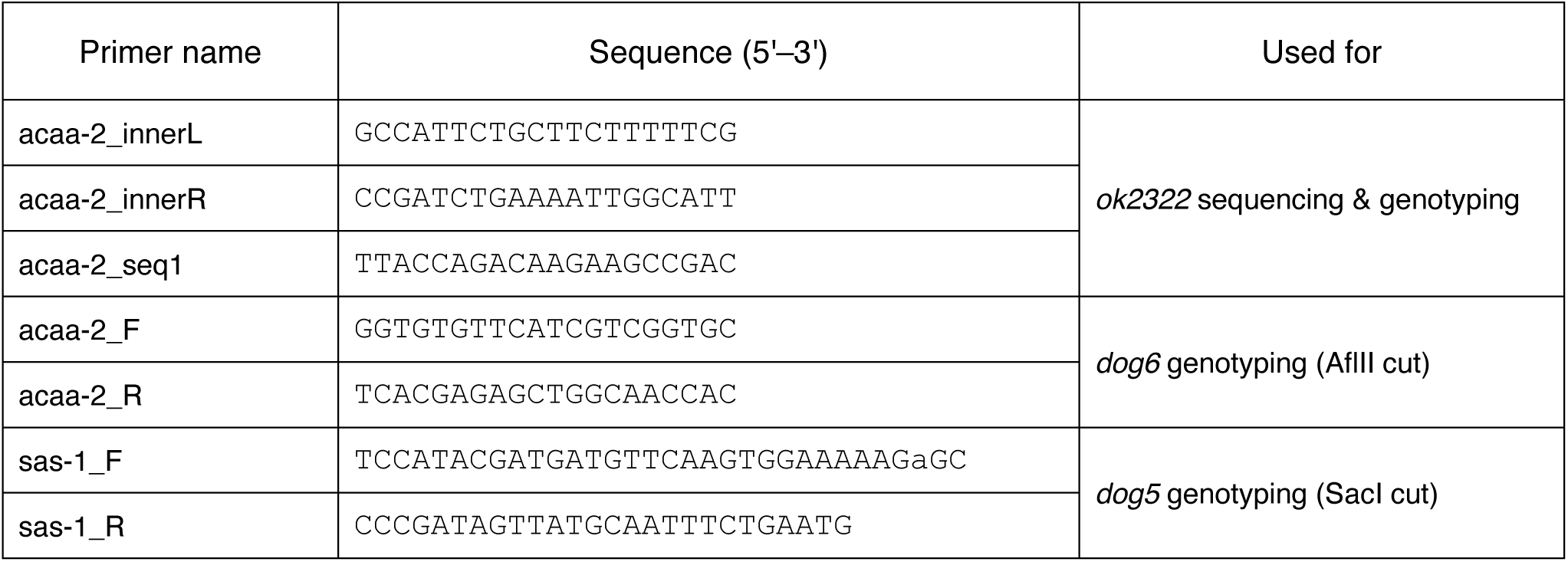
Primers used in this study.

## Movies

**Movie 1**

Representative three-fold stage embryo from wild-type (N2) mothers reared on *E. coli* HB101. Bright-field recording, 10 s.

**Movie 2**

Representative three-fold stage embryo from *acaa-2(dog6)* mothers reared on *E. coli* HB101. Bright-field recording, 10 s.

**Movie 3**

Representative three-fold stage embryo from wild-type (N2) mothers reared on *B. megaterium*DA1880. Bright-field recording, 10 s.

**Movie 4**

Representative three-fold stage embryo from *acaa-2(dog6)* mothers reared on *B. megaterium* DA1880. Bright-field recording, 10 s.

